# Herbarium specimens reveal herbivory patterns across the genus *Cucurbita*

**DOI:** 10.1101/2021.07.21.452357

**Authors:** Laura A. Jenny, Lori R. Shapiro, Charles C. Davis, T. Jonathan Davies, Naomi E. Pierce, Emily Meineke

**Affiliations:** Department of Organismic and Evolutionary Biology, Harvard University, Cambridge, MA; Department of Applied Ecology, North Carolina State University, Raleigh, NC; Harvard University Herbaria, Department of Organismal and Evolutionary Biology, Harvard University, Cambridge, MA; Department of Department of Ecology and Evolutionary Biology, University of British Columbia, Vancouver; African Centre for DNA Barcoding, University of Johannesburg; Department of Entomology and Nematology, University of California, Davis, Davis, CA

**Keywords:** *Cucurbita*, pumpkin, squash, herbivory, co-evolution, cucumber beetle, *Acalymma*, herbarium, *Erwinia tracheiphila*, plant-herbivore interactions

## Abstract

**PREMISE:** Quantifying how closely related plant species differ in susceptibility to insect herbivory is important for our understanding of variation in plant-insect ecological interactions and evolutionary pressures on plant functional traits. However, empirically measuring *in situ* variation in herbivory over the entire geographic range where a plant-insect complex occurs is logistically difficult. Recently, new methods have been developed to use herbarium specimens to investigate patterns in plant-insect interactions across geographic areas, and during periods of accelerating anthropogenic change. Such investigations can provide insights into changes in herbivory intensity and phenology in plants that are of ecological and agricultural importance.

**METHODS:** Here, we analyze 274 pressed herbarium samples from all 14 species in the economically important plant genus *Cucurbita* (Cucurbitaceae) to investigate variation in herbivory damage. This collection is comprised of specimens of wild, undomesticated *Cucurbita* that were collected from across their native range in the Neotropics and subtropics, and *Cucurbita* cultivars that were collected from both within their native range and from locations where they have been introduced for agriculture in temperate Eastern North America.

**RESULTS:** We find that herbivory is common on individuals of all *Cucurbita* species collected from throughout their geographic ranges; however, estimates of herbivory varied considerably among individuals, with greater damage observed in specimens collected from unmanaged habitat. We also find evidence that mesophytic species accrue more insect damage than xerophytic species.

**CONCLUSIONS:** Our study demonstrates that herbarium specimens are a useful resource for understanding ecological interactions between domesticated crop plants and co-evolved insect herbivores.

## INTRODUCTION

Herbaria were originally envisioned as archives to store samples for research related to plant morphology and taxonomy, later expanding to studies of phenology, phylogenetics and biogeography. For centuries, the use of herbarium specimens was largely restricted to studies on these topics. Only recently have methods been developed to also gather ecological information from herbaria samples. Studies using herbarium specimens have inferred human influences on plant ecological interactions with insect herbivores, insect pollinators, and microbial associates (Daru et al., 2018; Lughadha et al., 2018; Meineke et al., 2018b; Meineke and Davies, 2018; Meineke et al., 2018a). These collections have also been used to quantify how the increasing global footprint of human activities is changing geographic ranges, population sizes, and species interactions of many plant species (Lughadha et al., 2018; Meineke and Davies, 2018).

Specimens of domesticated crop plants and their wild relatives from herbaria may contain ecological information for species that are of particular importance to human food systems, although this remains largely unexplored. The plant genus *Cucurbita* (Cucurbitaceae) is an ecologically and agriculturally important model for understanding plant-insect interactions (Metcalf, 1979; Metcalf and Lampman, 1989; Shapiro and Mauck, 2018). Most experimental work on plant-insect ecological interactions involving *Cucurbita* has been conducted with cultivated populations in the introduced range and is notably concentrated in the Midwestern and Northeastern United States (Sasu et al., 2009; Sasu et al., 2010; Shapiro and Mauck, 2018). Very little is known about variation in plant-insect ecological interactions among *Cucurbita* species in both wild and cultivated settings, and across its native range in the American tropics and subtropics (Kates et al., 2017). The 14 species that comprise *Cucurbita* fall into two phylogenetic groups. There are six xerophytic (dry-adapted) perennial species native to arid areas in the Southwestern United States and Northwestern Mexico, and eight species of mesophytic (neither dry nor wet adapted) annuals native to an area spanning the southeastern United States through South America (Nee, 1990; Kates et al., 2017). Five mesophytic species have been domesticated for agriculture, and remain ecologically and economically important throughout their native range (Nee, 1990; Sanjur et al., 2002; Piperno and Stothert, 2003; Kates et al., 2017). Recent phylogenomic dating using 44 nuclear markers found that the xerophytic species are the ancestral group, and the mesophytic *Cucurbita* are the result of a radiation that occurred ~7 million years ago (Agrawal and Fishbein, 2008; Schaefer et al., 2009; Kates et al., 2017).

All *Cucurbita* spp. produce anti-herbivory secondary metabolites called ‘cucurbitacins’, which are among the most bitter compounds characterized. Cucurbitacins act as effective deterrents for nearly all insect and mammalian herbivores (Chambliss and Jones, 1966; Metcalf, 1979; Metcalf and Lampman, 1989), except for a small subset of highly co-evolved leaf beetles in the genus *Acalymma* (Coleoptera: Chrysomelidae: Luperini: Diabroticina). These beetles are among the only animals that can detoxify cucurbitacins and consume *Cucurbita* tissues, and all ~70 *Acalymma* spp. are obligately dependent on *Cucurbita* host plants in all life stages. For these beetles, cucurbitacins act as arrestants and feeding stimulants (Barber, 1946; Munroe and Smith, 1980; Samuelson, 1994; McCloud et al., 1995; Eben and Gamez-Virues, 2007; Gillespie et al., 2008; Eben and Espinosa de los Monteros, 2013). The ability of *Acalymma* spp. to metabolize cucurbitacins, and the obligate dependence of *Acalymma* beetles on *Cucurbita* host plants suggests this group of beetles have likely been an important selective pressure on *Cucurbita* over the estimated ~4 million years that they have been co-evolving (Metcalf, 1979; Metcalf and Lampman, 1989; Andrews et al., 2007; Gillespie et al., 2008; Eben and Espinosa de los Monteros, 2013). While some *Diabrotica* and *Epilachna* species also consume *Cucurbita* tissue, these beetles are polyphagous and feed on many other plant species (Carrol and Hoffman, 1980; Eben and Gamez-Virues, 2007), and the vast majority of chewing herbivory on *Cucurbita* is by *Acalymma* beetles (Du et al., 2008; Hladun and Adler, 2009).

Despite the economic and ecological importance of *Cucurbita,* variation in herbivory between species and cultivars, and over time and space is poorly understood. For example, one puzzling trait of the *Cucurbita-Acalymma* co-evolutionary complex is that *A. vittatum* transmits the fatal bacterial wilt, *Erwinia tracheiphila,* (Shapiro et al., 2014) but only in temperate Eastern North America. *Erwinia tracheiphila* does not occur in South America or Mesoamerica, which is the evolutionary center of origin and species radiations for both *Cucurbita* and *Acalymma* (Shapiro et al., 2016; Shapiro et al., 2018). The relative paucity of knowledge about plant-insect interactions throughout the native range of both partners constrains our ability to fully understand why this pathogen has such a restricted distribution, and whether other host plant populations may be at risk. It is possible that variation in herbivory may be one of the factors affecting *E. tracheiphila’*s current distribution.

Here, we quantify foliar beetle herbivory on the collection of all *Cucurbita* spp. at the Harvard University Herbaria, which houses *Cucurbita* specimens collected throughout the Neotropics and subtropics where *Cucurbita* are native, and in temperate parts of the Americas where cultivated *Cucurbita* have been introduced for agriculture. We use these data to contrast patterns of herbivory on domesticated *Cucurbita* versus wild relatives, and to examine how herbivory varies spatiotemporally. We find that specimens from mesophytic taxa have higher levels of herbivory damage compared to xerophytic specimens, and that specimens collected from the wild have higher levels of herbivory damage than domesticated specimens collected from managed gardens. This study also establishes a ‘proof of principle’ that herbarium specimens can be used to inform investigations into how human activities related to agriculture may be altering plant-insect ecological interactions that are relevant to agricultural production and food security.

## MATERIALS AND METHODS

### Specimen overview

We quantified herbivory on all 274 *Cucurbita* samples in the Harvard University Herbaria and recorded the label metadata for each specimen (Supplemental File 1). For some samples, species taxonomy has changed since the original collection. In these cases, the current taxonomy following Kates *et al.* (Kates et al., 2017) was applied. The original taxonomic assignments given by the collector, and the updated taxonomic assignments are both provided in Supplemental File 1. In other instances, label data were incomplete for location or date collected. For example, most of the samples were collected before handheld GPS units were widely available, so the location is often given relative to local roads, rivers, or other landmarks. In these cases, latitude and longitude information was approximated based on the location information provided. Location information was used to label samples as being in the geographic region where bacterial wilt disease is present (Northeastern and Midwestern North America) or outside of the region where the disease is present. Specimens were also coded to record whether they were likely collected from a human-managed garden or from a wild, unmanaged population. All samples collected from temperate regions were assigned as ‘garden’ because *Cucurbita* do not naturally occur in temperate climate zones, although we recognize that some of the ‘garden’ collections could be cultivated varieties that escaped agricultural settings and are growing as volunteers.

### Herbivory quantification

For all 274 samples, foliar herbivory was quantified following protocols described in Meineke *et al.* (Meineke et al., 2018b). Foliar herbivory damage was quantified using a grid matching the size of standard herbaria sheets (41centimeter by 25 centimeter) with 40 numbered five-centimeter by five-centimeter boxes. Random numbers were used to select five boxes within the 40-cell grid that contained some foliage. The foliage within each of the five randomly selected boxes was visually inspected using a dissecting microscope (10X magnification) for the presence or absence of herbivory damage. Evidence of feeding damage was recorded as a binary (presence/absence) trait for each of the five boxes.

Chewing damage from herbivores with mandibles presents as jagged or smooth-edged holes that measure between one and five millimeters in diameter and destroys both the mesophyll and the epidermis. Chewing damage from *Acalymma* leaf beetles often produces a pattern of small holes in the leaf because they avoid consuming the heavily lignified vasculature tissue around the xylem and phloem tubes (a pattern of damage referred to as ‘skeletonization’). In rare cases, we also found apparent damage from leaf miners, which is characterized by a thin (approximately one millimeter) path of dead epidermis cells signifying that herbivores have consumed the mesophyll but not the epidermis. We scored chewing and mining herbivory separately as follows: the total amount of foliar herbivory damage was rated on a scale of zero to five, where a score of zero means no boxes showed herbivory damage, and a score of five means all boxes showed herbivory damage. This scoring system resulted in two separate scores from zero to five for chewing and leaf mining damage per specimen.

One challenge in quantifying herbivory on herbarium specimens is distinguishing between natural herbivory (pre-collection) and herbivory damage that the samples received during storage (post-collection). In a previous study, Meineke *et al.* (Meineke et al., 2018a) found that pre-collection herbivory on the leaves of some plant taxa can be differentiated by the presence of a thin, darkened outline around the damaged area, indicating the plant was still alive when the herbivory caused localized cell death. Post-collection herbivory or storage-related damage is inferred if localized cell death does not occur around the damaged area (Meineke et al., 2018a). We found the same pre- vs. post-collection leaf damage morphologies on *Cucurbita* and therefore applied these same methods for distinguishing pre-collection herbivory. However, we could not assess herbivory on *Cucurbita* flowers, even though beetles also consume floral tissues (Anderson and Metcalf, 1986; Anderson, 1987; Metcalf and Lampman, 1990; Shapiro et al., 2012), because it was not possible to differentiate pre-collection insect herbivory from post-collection damage on delicate *Cucurbita* flowers.

### Statistical Analyses

Chewing damage was present on 183 (67%) of the total 274 specimens. Only two specimens displayed mining damage. Because there was so little damage from mining herbivores and leaf mining insects very rarely attack *Cucurbita,* only the damage from chewing herbivores was included in our statistical analyses.

We built a series of Bayesian models with the BRMS package in R (Gelman et al., 2015; Burkner, 2017, 2018) to explore the effects of time, space and environmental conditions on insect herbivory. In all models, the response was the total number of boxes with herbivory standardized by the total number of boxes scored (five boxes, see above). Effects of each predictor were estimated as the mean and 95% credibility intervals from posterior distributions. Predictor variables were scaled and centered at zero, and a predictor variable with a mean and 95% credibility interval (CI) that did not include zero was considered statistically important. A zero-inflated binomial error structure with default priors was specified for all models. The models were defined as:

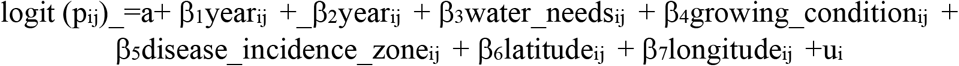

where grid cells with chewing damage is the number of grid cells with chewing damage by leaf beetle herbivores p on specimen i from species j, and n is a constant representing the number of grid cells examined on each specimen. We model logit(p_ij_) as a function of a, the intercept, year collected, species water needs (xerophytic or mesophytic), the growing condition (garden vs wild), the disease incidence zone, domestication status, latitude, longitude, and ui which is a grouping factor (random effects) of phylogenetic position. Rhat values of 1 indicated that all models converged.

Our main model includes all specimens (*N* = 274) (Model 1, Appendix S1). Phylogenetic relatedness among *Cucurbita* species was accounted for using a correlation matrix built from the genus phylogeny from Kates *et al.* (Kates et al., 2017) (Appendix S2) and the methods outlined in Turcotte *et al.* (Turcotte et al., 2014). Phylogenetic effects in the fitted model were estimated as the intra-class correlation (equivalent to Lynch’s lambda (Lynch, 1991)) using the “hypothesis” function in *brms*. We then created subsets of the data to test for effects of the same predictors described above; a description of the additional models including data subsets can be found in Appendix S1.

## RESULTS

### Specimen Distribution

Collection dates for *Cucurbita* specimens in the Harvard University Herbaria span 181 years, from 1835 through 2016. Sample location spans Southern Argentina to the Northeastern United States and includes three specimens collected in Caribbean islands (Fig 1, Table 1). Most samples (189 out of 274) were collected from Central and North America (north of the Panama Canal), and only 64 samples were collected from South America (south of the Panama Canal). The lower number of specimens from South America, where several *Cucurbita* species originate and remain culturally and economically important (Sanjur et al., 2002) (Piperno and Stothert, 2003), possibly reflects a bias towards collecting in areas closer to the Harvard University Herbaria (Daru et al., 2017).

**FIGURE 1:**
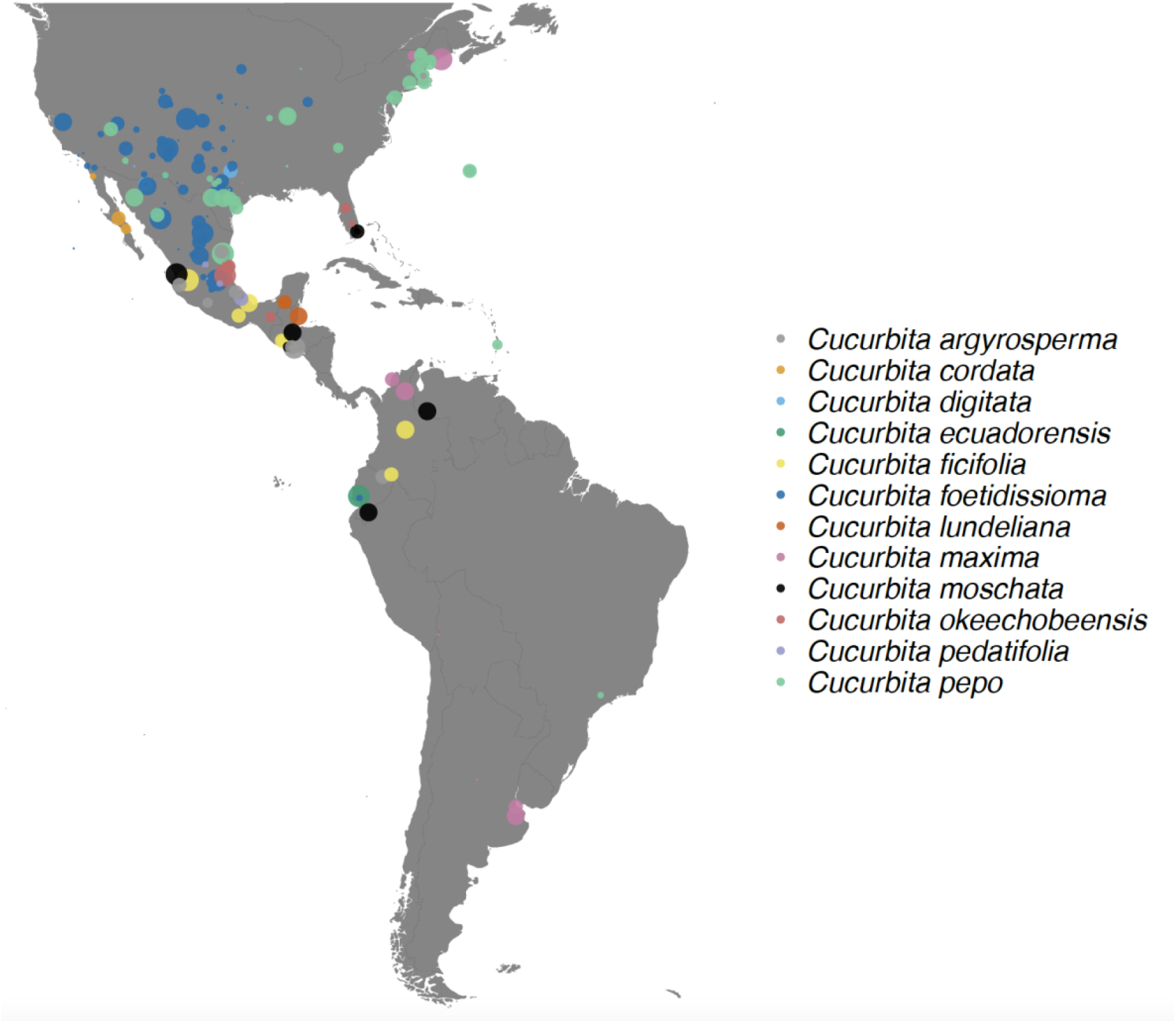
Map of the geographic distribution of *Cucurbita* specimens in the Harvard University Herbaria Collection with corresponding levels of herbivory damage shows that herbivory exists throughout the geographic range of the genera in the Americas. Different colors correspond to different *Cucurbita* species and larger sized points indicate more severe herbivory on the specimen.

**TABLE 1:** Specimens of *Cucurbita* in the Harvard University Herbaria span the time period from 1835-2016 and geographic locations in North, Central and South America.

| Current Species Taxonomy | Total Number of Samples | Date of First Sample | Date of Last Sample | Countries of Origin | Water Needs |
| --- | --- | --- | --- | --- | --- |
| *Cucurbita argyrosperma* | 23 | 1904 | 1991 | El Salvador, Mexico, United States | Mesophytic/Xerophytic |
| *Cucurbita cordata* | 10 | 1931 | 1974 | Mexico | Xerophytic |
| *Cucurbita digitata* | 8 | 1874 | 1965 | Mexico, United States | Xerophytic |
| *Cucurbita ecuadorensis* | 2 | 1913 | 1923 | Ecuador, United States | Mesophytic |
| *Cucurbita ficifolia* | 8 | 1886 | 1967 | Colombia, Guatemala, Mexico | Mesophytic |
| *Cucurbita foetidissima* | 79 | 1846 | 1993 | Mexico, United States | Xerophytic |
| *Cucurbita lundeliana* | 4 | 1931 | 1965 | Mexico, United States | Mesophytic |
| *Cucurbita maxima* | 19 | 1875 | 2016 | Argentina, Colombia, United States | Mesophytic |
| *Cucurbita moschata* | 12 | 1904 | 1993 | Colombia, Ecuador, El Salvador, Guatemala, United States | Mesophytic |
| *Cucurbita okeechobeensis* | 8 | 1858 | 1985 | Mexico, United States | Mesophytic |
| *Cucurbita pedatifolia* | 4 | 1890 | 1967 | Mexico | Xerophytic |
| *Cucurbita palmata* | 21 | 1875 | 1994 | Mexico, United States | Xerophytic |
| *Cucurbita pepo* | 76 | 1835 | 2016 | Brazil, Bermuda, Mexico, St. Lucia, United States | Mesophytic |
| TOTAL: | 274 | 1835 | 2016 |  |  |

Out of the 274 total samples, 62 were domesticated plants grown in a garden setting. Of the garden samples, 49 were collected in temperate Northeastern North America, where wild *Cucurbita* does not naturally occur, but domesticated varieties have been introduced for agriculture. The remaining 13 garden samples were collected in the American tropics and subtropics, where undomesticated *Cucurbita* wild relatives co-occur with domesticates. The three Caribbean samples were all *C. pepo* collected from gardens.

The two most abundant species in the collection were *C. foetidissima* and *C. pepo*, which both had between 70-80 samples (Fig 1). *Cucurbita ecuadorensis* was the least common species in the collection, with only 2 specimens. Seven out of 8 total specimens identified as *C. okeechobeenesis okeechobeensis* were collected from the eastern coast of Mexico, but these specimens are most likely misidentified because this species is rare, endangered and endemic to Florida, USA (Kates, 2019). This misidentification suggests some taxonomic uncertainty in species identification. *Cucurbita* species are all closely related, and some mesophytic species do not have diagnostic foliar morphological characteristics. Many samples also lack floral reproductive tissues that could provide more definitive taxonomic assignments (Chomicki and Renner, 2015). In these cases, only the use of high-resolution molecular markers will likely help in correctly determining the taxonomy assignments of the dried herbaria samples (Agrawal and Fishbein, 2008; Chomicki and Renner, 2015).

### Statistical Modeling

We found that chewing herbivory was common on *Cucurbita* specimens across all species examined. Bayesian R^2^ values ranged from 0.03-0.31 (Appendix S3). In the model including all species, herbivory did not vary over the 181-year timespan (Appendix S1, Model 1; Appendix S4), or between the disease incidence regions (Fig 2). Latitude and longitude were not significant predictors of herbivory damage, suggesting that insect herbivory is common throughout the *Cucurbita* geographic range where wild and domesticated genotypes occur. Mesophytic species had more chewing damage than xerophytic species, and no individual species drove this trend (Figs 2, 3). In addition, wild-collected specimens displayed more herbivory than specimens collected from gardens (Figs 2, 4). We also found a small, but detectable, relationship between phylogenetic relatedness and herbivory intensity (Appendix S5).

**FIGURE 2:**
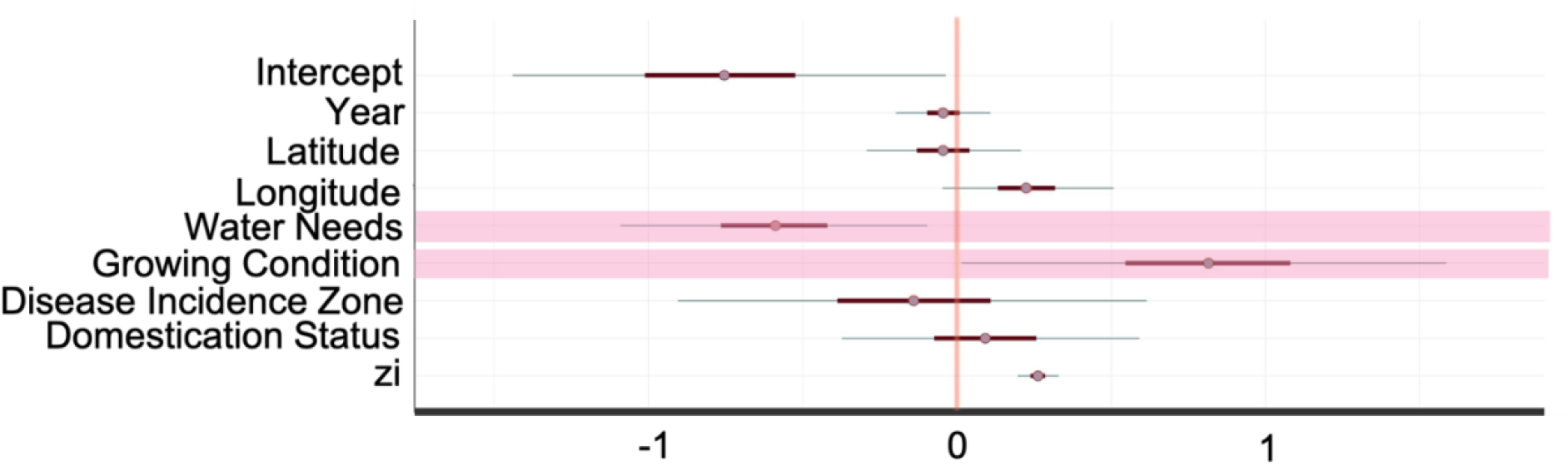
Model estimates showing the effects of time, space, plant characteristics, and environmental variables on herbivory damage to *Cucurbita* plants. Bold lines represent 80% credibility intervals and narrow lines represent 95% credibility intervals. Shading highlights the interaction term between herbivory damage and the water needs of the plant specimens (xerophytic vs mesophytic) and the growing conditions of the plant specimens (wild vs. garden).

**FIGURE 3:**
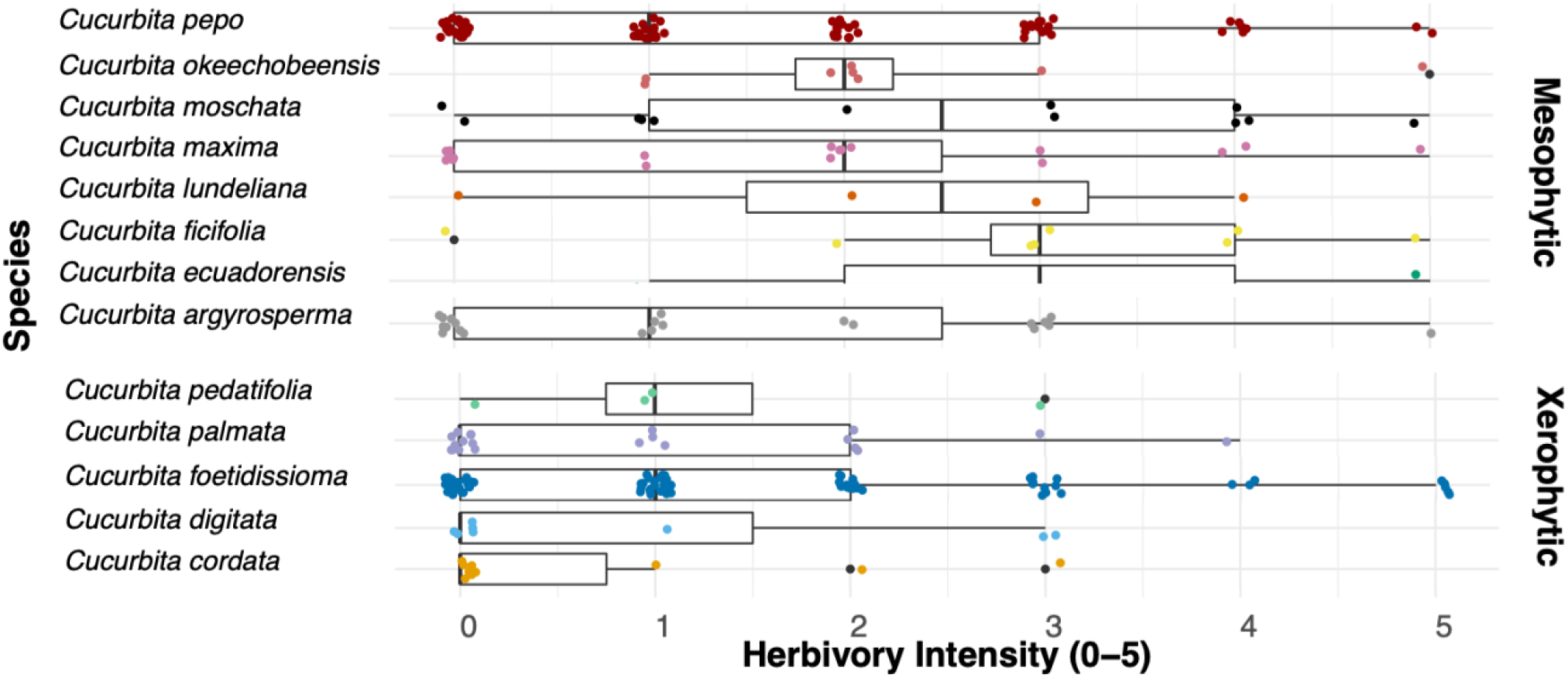
Mesophytic species of have significantly higher levels of herbivory damage compared to xerophytic species. Individual dots are colored according to species and represent individual samples in the Harvard University Herbaria Collection were scored on a scale of 0-5 where zero indicates little herbivory damage and five represents the highest level of herbivory damage.

**FIGURE 4:**
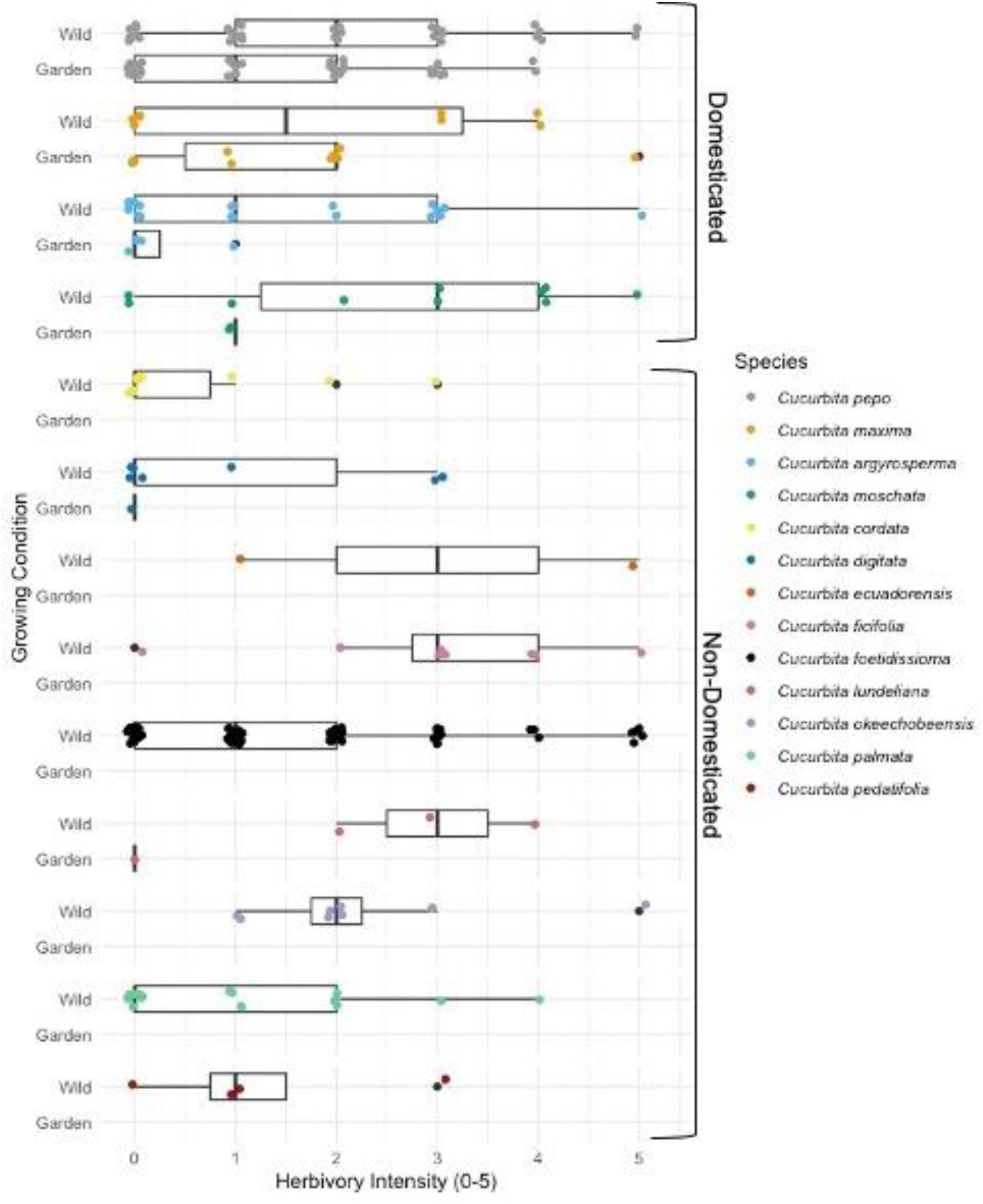
Specimens collected from wild settings displayed more herbivory than specimens collected from gardens. Individual dots are colored according to species and grouped into domesticated and non-domesticated species types. Herbivory intensity was scored on a scale of 0-5 for each specimen where zero indicates little herbivory damage and five represents the highest level of herbivory damage.

## DISCUSSION

Herbivory by mandibulate, chewing herbivores was common on herbarium specimens from all *Cucurbita* species, and across specimens from this genus gathered throughout tropical, subtropical and temperate America over a 181-year collection period. We found that mesophytic *Cucurbita* accrued more herbivory damage than xerophytic species, and specimens collected from the wild experienced more herbivory than those grown under cultivation. Our results suggest that anthropogenic changes, perhaps from the domestication process or host plant movement, have influenced plant-insect herbivore interactions within this group of agricultural food plants.

*Cucurbita* species are ancestral to xeric habitats in northwestern Mexico and the Southwestern part of the United States; mesophytic species evolved as a result of radiation into wetter habitats throughout the American tropics and subtropics ~7 million years ago (Hurd et al., 1971; Schaefer et al., 2009; Kates et al., 2017). Our finding that *Cucurbita* specimens from mesophytic habitats have higher levels of herbivory damage suggests this evolutionary transition from dry to moderate habitats may have affected *Cucurbita*-herbivore interactions. Evolutionary transitions to new habitats can provide many advantages to plants, including the possibility of escaping from herbivores (Agrawal, 2008). However, contrary to this hypothesis, we find that mesophytic *Cucurbita* species display herbivory damage than xerophytic species. This suggests that, while *Cucurbita* have spread successfully throughout subtropical regions in the Americas, escape from herbivory was not likely a facilitating factor in this range expansion (Bang and Faeth, 2011).

We also find evidence that cultivation affects herbivory patterns in *Cucurbita.* Five mesophytic species were domesticated for agriculture within the past ~10,000 years (Kates et al., 2017). The finding that herbivory is higher in wild specimens than in garden-collected specimens suggests two non-mutually exclusive mechanisms. First, it is possible that domesticates are less damaged by co-evolved herbivores because crop breeding has reduced the amount of cucurbitacins in domesticated cultivars. Rather than being deterred by defensive compounds like most herbivores, the co-evolved beetles that feed on *Cucurbita* are attracted to cucurbitacins and selectively feed on them, and *Cucurbita* have evolved to tolerate some amount of herbivory from the few co-evolved leaf beetle herbivores that are able to consume foliage containing cucurbitacins (Agrawal and Fishbein, 2008) (Strauss and Agrawal, 1999). *Cucurbita* plants grown in gardens may have lower cucurbitacin levels as a result of the process of domestication for agriculture (Brzozowski et al., 2019), reducing herbivory pressure from leaf beetle herbivores. Alternatively, plants grown in gardens may be attacked less simply because of anthropogenic interventions such as insecticides or lower general pest pressure in anthropogenic habitats. Because our dataset could not distinguish cultivated samples grown in gardens or agricultural fields from domesticates that are growing as weeds, experiments testing beetle attraction to, and herbivory on wild vs. domesticated *Cucurbita* will be necessary to distinguish these two hypotheses.

We also investigated the extent to which patterns in herbivory may co-vary with the incidence of cucurbit bacterial wilt caused by *Erwinia tracheiphila.* Our results indicating that herbivory by co-evolved beetles is ubiquitous throughout the Americas, including in regions outside where the pathogen occurs, supports previous investigations that found agricultural intensification and crop plant introductions, and not the geographic distribution of the beetle vector, underlie the recent emergence of this pathogen (Shapiro et al., 2016; Shapiro et al., 2018). Our finding that herbivory is ubiquitous throughout the time and geographic locations surveyed provides evidence that feeding frequency from obligate beetle vectors does not restrict the geographic distribution of the disease.

## CONCLUSION

While there is immense value in using herbaria specimens for describing plant-biotic ecological interactions, quantifying herbivory on herbarium specimens also presents challenges. *Cucurbita* species, in particular, are morphologically similar and difficult to identify. The species delimitations and taxonomy of *Cucurbita* have changed several times during the last several decades, and it is possible that some specimens in the Harvard University Herbaria collection have been assigned names based on outdated nomenclature. Time and funding permitting, additional data using molecular barcode markers would be valuable for classifying specimens. Nonetheless, our study demonstrates that herbarium specimens represent a rich source of species interactions data that can provide unique insights spanning an entire, widely distributed plant genus for which herbivory data are sparse across space and time.

## Acknowledgments

LJ was supported by a Myvanwy M. and George M. Dick Scholarship Fund for Science Students from Harvard Organismic and Evolutionary Biology, a Program for Research In Science and Engineering (PRISE) Fellowship, and a Harvard Microbial Sciences Initiative (MSI) Fellowship. This material is based on work supported by the National Science Foundation Postdoctoral Research Fellowship in Biology to EKM under grant no. 1611880. EKM was also supported by the UC Davis Department of Entomology and Nematology.

## Author contributions

LJ, LS, and EM conceived of the study. LJ performed the data collection of herbarium specimen. LJ, EM, CD, and JD developed analytical methods. LJ and EM analyzed data and all authors interpreted data. LJ, LS, EM wrote the first draft of the manuscript, and all authors revised the manuscript.

## Data availability

All data is contained in the Github repository: https://github.com/laurajenny/Cucurbita_herbaria_sup.git

## Supporting Information

Additional supporting information may be found online in the Supporting Information section at the end of the article.

Appendix S1: Bayesian modeling descriptions and results

Appendix S2: Correlation matrix file for phylogeny of *Cucurbita*

Appendix S3: R-squared values for Bayesian models

Appendix S4: Graph of herbivory on all specimens over 181-year timespan

Appendix S5: Phylogenic tree of species relatedness and herbivory intensity

